# Plasma lipid and liporotein biomarkers in LBC1936: Do they predict general cognitive ability and brain structure?

**DOI:** 10.1101/2020.07.09.194688

**Authors:** Sarah E. Harris, Stuart J Ritchie, Gonçalo D S Correia, Beatriz Jiménez, Chloe Fawns-Ritchie, Alison Pattie, Janie Corley, Susana Muñoz Maniega, Maria Valdés Hernández, John M. Starr, Derek Hill, Paul Wren, Mark E. Bastin, Matthew R Lewis, Joanna M. Wardlaw, Ian J. Deary

**Affiliations:** Lothian Birth Cohorts group, Department of Psychology, University of Edinburgh, 7 George Square, Edinburgh EH8 9JZ, UK; Social, Genetic, and Developmental Psychiatry Centre, King’s College, London SE5 8AF, UK; National Phenome Centre, Department of Metabolism, Digestion and Reproduction, Imperial College London SW7 2AZ, UK; Brain Research Imaging Centre, Neuroimaging Sciences, The University of Edinburgh, Chancellor’s Building, 49 Little France Crescent, Edinburgh EH16 4SB, UK; UK Dementia Research Institute at the University of Edinburgh, Edinburgh BioQuarter, Edinburgh EH16 4SB, UK; Alzheimer Scotland Dementia Research Centre, University of Edinburgh, 7 George Square, Edinburgh EH8 9JZ, UK; Department of Medical Physics and Biomedical Engineering, University College London WC1E 6BT, UK; ESCAPE Bio, San Francisco, CA 94080, USA; Scottish Imaging Network, A Platform for Scientific Excellence (SINAPSE) Collaboration, 300 Bath St, Glasgow G2 4LH, UK

## Abstract

Identifying predictors of cognitive ability and brain structure in later life is an important step towards understanding the mechanisms leading to cognitive decline and dementia. This study used ultra-performance liquid chromatography mass spectrometry (UPLC-MS) and nuclear magnetic resonance (NMR) to measure targeted and untargeted metabolites, mainly lipids and lipoproteins, in ∼600 members of the Lothian Birth Cohort 1936 (LBC1936) at aged ∼73 years. Penalized regression models (LASSO) were then used to identify sets of metabolites that predict variation in general cognitive ability and structural brain variables. UPLC-MS-POS measured lipids, together predicted 19% of the variance in total brain volume and 17% of the variance in both grey matter and normal appearing white matter volumes. Multiple subclasses of lipids were included in the predictor, but the best performing lipid was the sphingomyelin SM(d18:2/14:0) which occurred in 100% of iterations of all three significant models. No metabolite set predicted cognitive ability, or white matter hyperintensities or connectivity. Future studies should concentrate on identifying specific lipids as potential cognitive and brain-structural biomarkers in older individuals.

## Introduction

Ageing populations have an increasingly greater proportion of individuals affected by cognitive decline and dementia. To further our understanding of these important medical and societal problems, it is important to identify metabolite predictors of individual differences in cognitive ability and brain structure in later life ^1–4^. The present study investigates one set of potential predictors of cognitive and brain-structural differences: plasma lipid and lipoprotein biomarkers.

Lipids are important metabolites that form essential components of cell membranes and are involved in cell signalling processes ^5^. Lipoproteins are responsible for transporting water-insoluble lipids in blood and also carry apolipoproteins which determine the structure and function of the lipoproteins. Plasma lipid levels can be influenced by genetic ^6^ and environmental factors, including diet ^7^, exercise ^8^ and medication ^9^. Dysregulation of lipids has been associated with poorer cognitive function, cognitive decline, Alzheimer’s disease (AD), other dementias, and white matter hyperintensities (WMH), although — as one might expect for a relatively new area of research — results are not consistent ^10^.

High total cholesterol, low-density lipoproteins, triglycerides and apolipoprotein B, and low high-density lipoproteins have been associated with poorer cognitive performance and increased risk for dementia, particularly vascular dementia, in some, but not all studies (reviewed in ^10^). Inconsistencies may be due to the age of the subjects, the cognitive domains tested, dementia diagnosis criteria, and the lipids measured. Sex specific and non-linear associations of serum lipid levels with cognitive function have also been identified in middle- to older-aged individuals ^11^ and a study in the Lothian Birth Cohort 1936 (LBC1936) indicated that associations with cognitive function in older age may be confounded by childhood cognitive function ^12^.

Cholesterol is an important lipid in the brain, vital for synapse formation and development, dendrite differentiation, axonal elongation, and long-term potentiation ^13^. The blood-brain barrier (BBB) separates brain metabolised cholesterol and lipoproteins from those in plasma. Increased BBB permeability is associated with cognitive decline and has been shown to occur in rats fed diets high in saturated fats and cholesterol ^14–16^. Anti-inflammatory and lipid-lowering agents have been shown to reverse this phenomenon ^14^. Increased HDL-cholesterol and decreased LDL-cholesterol were associated with increased WMH in a longitudinal study of 1919 individuals aged over 65 years ^17^ and hyperlipidemia was associated with less severe WMH in patients with acute ischaemic stroke in two independent cohorts (N=1135) ^18^, suggesting that hyperlipidemia may actually protect against small vessel disease. Lower HDL-cholesterol predicted increased WMH volume from age 73 to 76 years in the LBC1936 ^19^.

Extracellular plaques consisting of amyloid beta peptides are one of the main neuropathologies associated with AD. Genetic variants in genes involved in the amyloid pathway are associated with AD. Apolipoprotein E (APOE) regulates transport of cholesterol and lipids and mediates clearance of plasma lipoproteins. The *APOE* e4 allele, that is associated with an increased risk of late-onset AD as well as risk of steeper non-AD cognitive decline ^20^, impairs clearance of the plasma lipoproteins, enhancing amyloid beta aggregation ^21^. Sphingolipids, including sphingomyelin and ceramide, have important functional and structural roles in cellular membranes. Sphingomyelin and cholesterol interact to influence membrane permeability ^22^.

Modern analytical technologies such as nuclear magnetic resonance (NMR) and ultra-performance liquid chromatography mass spectrometry (UPLC-MS) enable the broadscale detection and measurement of chemical constituents within biofluids and tissue extracts. Although both techniques have been used extensively to measure small molecule metabolites, NMR has emerged in recent years as a popular and reproducible means to quantify lipoprotein species ^23^. Similarly, UPLC-MS is now commonly applied for the measurement of complex lipids from human biofluids and tissue extracts. Such analyses may be targeted to measure sets of well-defined species or untargeted whereby thousands of lipid-related features are detected in a single analysis. In the latter case, those measurements showing a positive association with a trait of interest require chemical annotation prior to downstream interpretation of the results.

Large scale metabolomics studies of cognitive function and decline are starting to identify biological mechanisms and biomarkers of cognitive impairment in later life ^24,25^. A recent study of lipidome evolution in mammalian tissues showed a significant excess of lipid concentration changes in the brain cortex, with cortical changes clustered in glycerolipid, glycerophospholipid, and linoleic acid metabolism pathways, all of which have previously been implicated in neurodegenerative disorders ^26^. Serum levels of a glycerophosphocoline (a subclass of glycerophospholipids) have been suggested as potential marker of visceral fat related peripheral inflammation, associated with changes in brain structure and function ^27^.

In the present study we used penalized regression models (LASSO) to identify sets of metabolites (principally lipids and lipoproteins) in both targeted and untargeted UPLC-MS and NMR spectroscopy data sets that might be used to predict some of the variation in general cognitive ability and structural brain variables in the LBC1936.

## Materials and Methods

### Lothian Birth Cohort 1936 (LBC1936)

The LBC1936 consists of 1091 individuals, most of whom, at the age of ∼11 years, took part in the Scottish Mental Survey of 1947, when they took a validated test of cognitive ability, the Moray House Test (MHT) version 12. Between 2004 and 2007, they were recruited to a study to determine influences on cognitive ageing at age ∼70 years and have now taken part in five waves of testing in later life (ages ∼70, ∼73, ∼76, ∼79 and ∼82 years). At each wave, they have undergone a series of cognitive and physical tests, with brain magnetic resonance imaging (MRI) introduced at age 73 years. For the present study plasma was extracted from lithium heparin collected blood at Wave 2 at a mean age of 72.5 (SD 0.7) years and stored at −80°C ^28^.

### Metabolomics

Plasma metabolic profiles were measured at the National Phenome Centre at Imperial College London. Plasma samples were shipped on dry ice and stored at −80°C upon receipt prior to aliquoting for subsequent analyses by thawing in batches of 80 overnight at 4°C, centrifuging, and decanting the homogenous supernatant into 96 well plates as previously described ^29^. Sample plates were stored at −80°C until needed for analysis. Quality control materials including a study reference (SR) sample (equal-parts pool of all study samples) were prepared as previously described ^29^.

#### Lipidomics

Plasma samples were prepared for lipidomics analysis by dilution with isopropanol^30^ as previously described for serum analysis ^29^ with the following modification: 100 µL of sample were thawed at 4°C for 2h and protein precipitation was achieved by addition of four parts of an isopropanol solution containing reference standards to one part plasma. UPLC-MS was performed using Acquity UPLC chromatography systems (Waters Corporation, Milford, MA, USA) and Xevo G2-S Q-ToF mass spectrometers with electrospray interface (Waters Corp., Manchester, UK). Reversed-phase chromatography was conducted using an Acquity 2.1×100mm BEH C8 column, and separate analyses were performed for positive and negative ion detection (UPLC-MS-POS and UPLC-MS-NEG respectively). Details of the chromatographic solvent system, gradient elution, mass spectrometer parameters, and data acquisition have been published previously ^29^. Targeted integration of signals arising from known lipid species in the UPLC-MS datasets was performed using the peakPantheR R package ^31^ and an in-house database of empirical retention time and theoretical m/z values for annotated lipids. The nPYc-Toolbox ^32^ was then used to correct ion intensities for sample analysis-order intensity drift using a LOWESS smoother estimated using the intensities measured across repeated injections of the SR. Following drift correction, the SR samples and their serial dilution series were used to calculate feature-wise coefficients of variation (CV) and Pearson correlation coefficients with the dilution factor, respectively. Only signals with a CV lower than 30% and a correlation with dilution higher than 0.7 were kept for statistical analysis.

#### Biocrates AbsoluteIDQ® p180

The Biocrates AbsoluteIDQ® p180 (Biocrates Life Sciences AG, Innsbruck, Austria) kit provided absolute quantification for 53 compounds, and semi-quantitative measurements for a further 135 compounds. Data were acquired on Waters TQ-S instruments (Waters Corp., Manchester, UK) using Acquity UPLC systems (Waters Corporation, Milford, MA, USA).

#### Nuclear Magnetic Resonance (NMR) Spectroscopy

An adaptation of the standard metabolic profiling method for NMR ^33^ was developed and validated for this analysis due to the limited sample volume available. Briefly, 40 µL of plasma were mixed with 40 µL of aqueous buffer containing 20% of D_2_O 75 mM of NaH2PO_4_, 4.6 mM of TSP (3-(trimethylsilyl)-2,2,3,3-tetradeuteropropionic acid or TMSP-d4) and 6.2 mM of NaN_3_ at pH 7.4 in an Eppendorf and vortexed. Subsequently, 60 µL of the mixture were transferred to a 4’’ 1.7 mm tube using a 100 µL Hamilton syringe. Samples were prepared in batches of 80 samples and subsequently split into two batches of 27 and one of 26 for analysis. One SR and one LTR samples were included in each of the batches with a total of 6 QCs included per 80 samples. An additional 600 µL of LTR sample was prepared and run during the calibration of each spectrometer in order to transfer the IVDr methods to the small volume samples. Each 27-sample batch was transferred to the refrigerated sample handling robot (SampleJet, Bruker Corporation, Germany) of a one of three Bruker Avance III HD 600 NMR spectrometers for analysis and parallel acquisition was completed within 24 hours of sample preparation. 1D NMR profiling was acquired using the 1D-NOESY presat pulse sequence, a spin-echo experiment using the 1D-CPMG (Carr-Purcell-Meiboom-Gill) presat pulse sequence and *J*-res 2D experiments were run in automation at 310 K using a BBI probe with z-gradients and high degree shimming. A relaxation edited spin-echo experiment, the CPMG sequence, is often acquired for plasma samples in order to filter out fast relaxing signals belonging to proteins and other large molecules in the ^1^H NMR spectrum, providing a better resolution for the slow relaxing signals belonging to metabolites and small molecules. 128 Free Induction Decays (FID) were accumulated for each experiment in 96 K points using a 30 ppm window centred at 4.78 ppm for the general profile while it was set to 20 ppm for the spin-echo, and 6 scans and 40 planes were acquired for the 2D *J*-resolved experiment. The relaxation delay was set at 4 s for the 1Ds while the *J*res was set to 2s and a water pre-saturation pulse was applied during this period to cancel the water signal.

Quality of the spectra were assessed following the criteria described in ^33^. Any samples not passing the QC criteria were rerun at least once, and otherwise left outside the final dataset. Data was prepared for statistical analysis with the nPYc-Toolbox ^32^, by re-calibrating the chemical shift scale using the α-glucose signal at δ 5.233 and re-interpolating all spectra to a common chemical shift axis.

#### B.I.-LISA platform

One hundred and five specific lipoproteins were measured using a modified version of the Bruker *B.I.-Lisa* platform (Bruker IVDr Lipoprotein Subclass Analysis). The platform is developed and optimised for the analysis of 300 μL aliquot plasma samples as described in^34^. In order to calculate the correction factor needed to extend the applicability of the method to the current analysis, an independent set of 622 samples was analysed both using the standard protocol and the reduced volume one used for this project. The error associated to the current measurements is increased with respect to the precision and accuracy reported in ^23^. The lipoproteins measured were cholesterol, free cholesterol, phospholipids, triglycerides, apolipoproteins A1, A2, B and particle numbers for the overall concentrations contained in plasma but also information on the concentration of the subclasses of those lipoproteins as obtained by ultracentrifugation was provided.

### Cognitive Tests

Cognitive ability was assessed using 13 cognitive tests taken by the LBC1936 sample, all at age 72.5 (SD 0.7) years (Wave 2). The tests were the Matrix Reasoning (MR), Block Design (BD), Digit Span Backward (DSB), Symbol Search (SS), and Digit-Symbol Substitution (DSS) tests from the Wechsler Adult Intelligence Scale, Third UK Edition ^35^; the Logical Memory (LM), Verbal Paired Associates (VPA), and Spatial Span (SSP) tests from the Wechsler Memory Scale, Third UK Edition ^36^; the National Adult Reading Test (NART) ^37^; the Wechsler Test of Adult Reading (WTAR) ^38^; a phonemic Verbal Fluency (VF) test ^39^; a Choice Reaction Time (CRT) test ^40^; and an Inspection Time test (IT) ^41^. Structural equation modeling was used to produce a latent variable from the variance shared among these tests in the following manner. On the basis of previous factor-analytic work in this sample ^42^, the tests were organized into a higher-order confirmatory factor-analytic model that included latent domains of visuospatial ability (indicated by the tests of MR, BD, and SSP), verbal memory (indicated by LM, VPA, and DSB), processing speed (indicated by SS, DSS, CRT, and IT), and crystallized ability (indicated by NART, WTAR, and VF), all of which were indicators of the highest-level latent variable, which we named “general cognitive ability”. From this model, the factor scores for the general cognitive ability factor were saved, and were used to index each individual participant’s general cognitive ability in our prediction models. Structural equation modelling was performed, and the latent variables were produced, using the lavaan package for R ^43^.

### Brain MRI

Whole-brain structural and diffusion tensor MRI data were acquired using a 1.5T GE Signa Horizon scanner (General Electric, Milwaukee, WI, USA) at the Brain Research Imaging Centre, the University of Edinburgh, shortly after cognitive testing and plasma collection. Mean age at scanning was 72.7 (SD 0.7) years. Full details are given in ^44^. Total brain, grey matter, normal appearing white matter and WMH volumes were calculated using a semi-automated multispectral fusion method, described previously ^45–47^.

The diffusion tensor MRI protocol employed a single-shot spin-echo echo-planar (EP) diffusion weighted sequence in which diffusion-weighted EP volumes (*b* = 1000 s mm^−2^) were acquired in 64 non-collinear directions, together with seven T2-weighted EP volumes (*b* = 0 s mm^−2^). This protocol was run with 72 contiguous axial slices with a field of view of 256 × 256 mm, an acquisition matrix of 128 × 128 and 2mm isotropic voxels ^47^.

White matter connectivity data were created using the BEDPOSTX/ProbTrackX algorithm in FSL (https://fsl.fmrib.ox.ac.uk) and 12 major tracts of interest were segmented using Tractor (https://www.tractor-mri.org.uk) scripts: the genu and splenium of the corpus callosum, and bilateral anterior thalamic radiations, cingulum bundles, uncinate, arcuate and inferior longitudinal fasciculi. Tract-average white matter fractional anisotropy (FA) and mean diffusivity (MD) were derived as the average of all voxels contained within the resultant tract maps. General factors of FA (*g*FA) and MD (*g*MD) were derived from a confirmatory factor analysis with a single general variable indicated by either FA or MD values from across all 12 tracts, to reflect the well-replicated phenomenon of common microstructural properties of brain white matter ^48–50^. As above, the confirmatory factor analytic models used the *lavaan* package.

Brain imaging data and metabolite measures were available for a maximum of 668 and 595 individuals, respectively.

## Statistical analyses

### LASSO prediction models

In order to produce metabolomic predictors of general cognitive function and of neuroimaging measures, Least Absolute Shrinkage and Selection Operator (LASSO) penalized regression models were used ^51^. These models produce a sparse solution to a prediction problem, dealing with a large number of predictors and the potential for collinearity among them. They do so by shrinking (penalizing) the coefficients of many of the predictors to zero in their regularisation process; in this sense, they also allow for variable selection, since a smaller (and thus more manageable) number of predictors might survive the process than at the outset.

We ran a series of LASSO models, attempting to predict variation in each brain MRI variable or cognitive outcome from all of the features derived from each metabolomic array in turn. For each model, the LBC1936 data were split at random into a training sample (80% of the data) and a testing sample (20% of the data). We first performed a LASSO regression in the training sample alone. Using 10-fold cross-validation, the penalization parameter (often denoted as λ) that provided the lowest prediction error in the training set was identified. The predicted values from a regression with this cross-validated penalization parameter were then taken forward to the testing sample, where they formed a predictor that was regressed on the relevant outcome variable (cognitive ability or brain volume). The R^2^ (proportion of variance accounted for by the predictor, expressed in the results below as a percentage) was recorded for each model. This procedure was repeated 1,000 times, with the LBC1936 dataset being split into a different random training and testing set each time. The mean R^2^ from the 1,000 iterations was calculated, and the standard deviation across all the iterations was used as the (bootstrapped) standard error for this mean value. Where the computational demands were too high for this iterative procedure to successfully be performed, the number of model iterations was reduced.

The LASSO models were run using the *glmnet* package for R ^52^, with wrapper functions from the *caret* package ^53^.

In a final feature-selection stage, metabolites that occurred in > 75% of the 1000 iterations were identified for those models where the metabolites explained a significant proportion of the variance in the outcome (that is, the confidence interval of their *R*^2^ value, derived using the bootstrapped standard error, did not cross zero). Since a large number of *R*^2^ values were calculated across all the arrays, we corrected their significance levels using the False Discovery Rate correction. This adjustment for multiple comparisons meant that, at the end of our analysis, only the strongest predictions would remain statistically significant.

### Sensitivity analyses

For models where the metabolites explained a significant proportion of the variance in the outcome, two sensitivity analyses were performed: 1) using rank-based inverse normal transformed metabolites (ranknorm() function from the R package *RNOmni* ^54^) to ensure that predictions were not being driven by outlying measurements; 2) correcting brain volumes for intracranial volume (a proxy measure of previous brain volume) to identify specific associations with age-related brain volume atrophy. Correlations between total brain, grey matter and normal appearing white matter volumes, and intracranial volume were also calculated.

## Results

Descriptive statistics for general cognitive ability and brain MRI variables are shown in Table 1. Eight hundred and sixty-six subjects provided cognitive data, and between 656 and 668 provided data on the brain imaging variables.

**Table 1.**
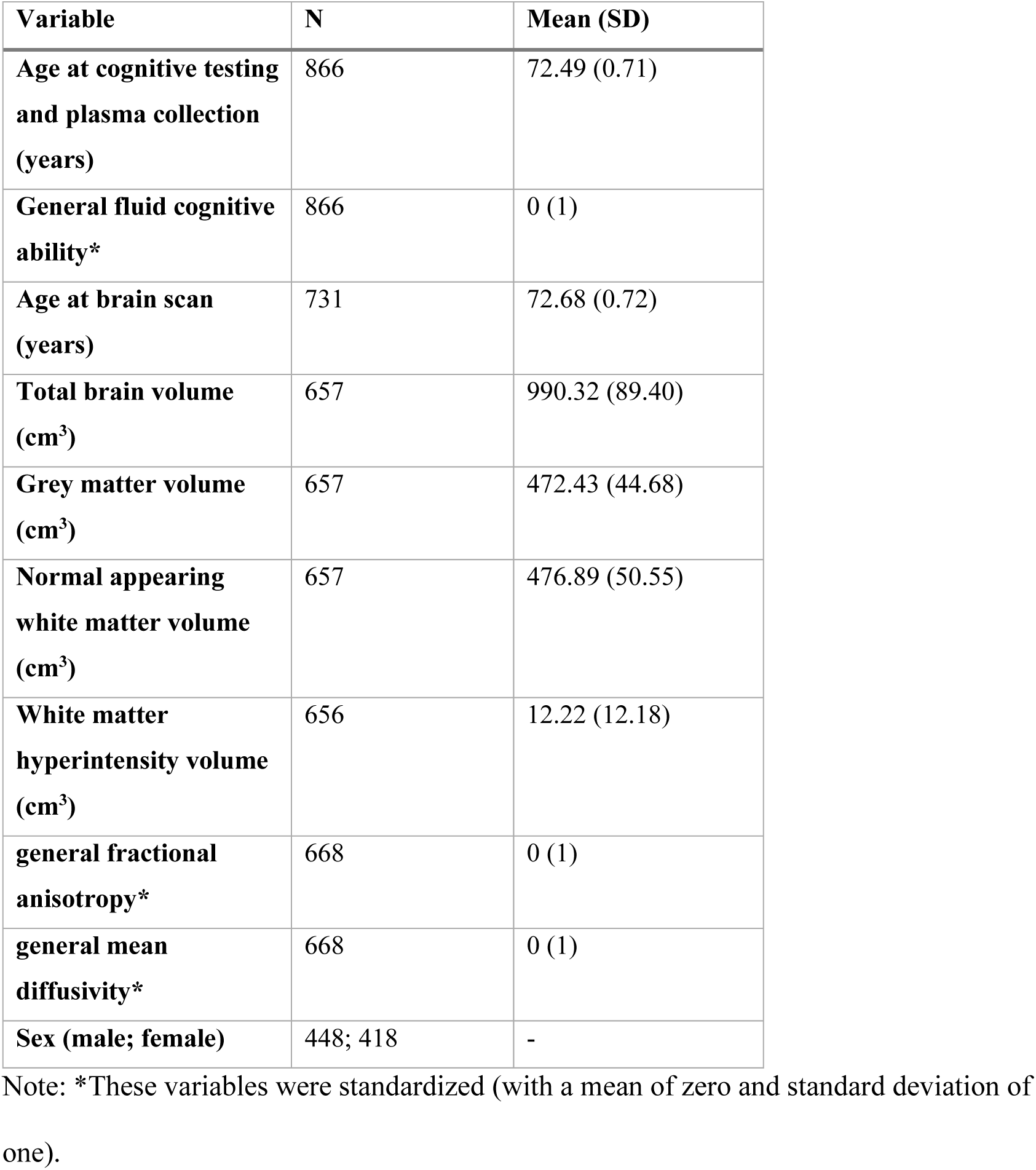
Summary descriptive data for LBC1936

| **Variable** | **N** | **Mean (SD)** |
| --- | --- | --- |
| **Age at cognitive testing and plasma collection (years)** | 866 | 72.49 (0.71) |
| **General fluid cognitive ability\*** | 866 | 0 (1) |
| **Age at brain scan (years)** | 731 | 72.68 (0.72) |
| **Total brain volume (cm<sup>3</sup>)** | 657 | 990.32 (89.40) |
| **Grey matter volume (cm<sup>3</sup>)** | 657 | 472.43 (44.68) |
| **Normal appearing white matter volume (cm<sup>3</sup>)** | 657 | 476.89 (50.55) |
| **White matter hyperintensity volume (cm<sup>3</sup>)** | 656 | 12.22 (12.18) |
| **general fractional anisotropy\*** | 668 | 0 (1) |
| **general mean diffusivity\*** | 668 | 0 (1) |
| **Sex (male; female)** | 448; 418 | - |

Note: \*These variables were standardized (with a mean of zero and standard deviation of one).

For the Biocrates AbsoluteIDQ^®^ p180 platform, 171 of 188 metabolites were detected in LBC1936 (Supplementary Table 1). All 105 B.I.-LISA platform metabolites were detected. (Supplementary Table 2). From the UPLC-MS features, 193 and 55 metabolites were annotated, respectively, in the UPLC-MS-POS and UPLC-MS-NEG datasets (Supplementary Tables 3 and 4). Untargeted NMR-NOESY and NMR-CPMG sequences each detected 18,646 features. Metabolites detected by each platform were used in platform-specific penalized regression models (LASSO) as described above.

Prediction results are displayed in Table 2. Only a small number of predictions, all from the UPLC-MS-POS array, remained statistically significant after correction for multiple comparisons. However, in those cases, the predicted variance from the LASSO models was substantial.

**Table 2.** Results from LASSO prediction models. *g* = general cognitive ability, TBV = total brain volume, GMV = grey matter volume, NAWM = normal appearing white matter volume, WMHV = white matter hyperintensity volume *g*FA = general fractional anisotrophy, *g*MD = general mean diffusivity. *500 iterations, ^#^100 iterations were run rather than 1000 due to high computational demands. FDR corrected significant findings are indicated in b

|  | Biocrates |  |  | B.I.-LISA |  |  | UPLC-MS-POS |  |  | UPLC-MS-NEG |  |  | NMR-NOESY |  |  | NMR-CPMG |  |  |
| --- | --- | --- | --- | --- | --- | --- | --- | --- | --- | --- | --- | --- | --- | --- | --- | --- | --- | --- |
|  | R<sup>2</sup> | SE | P | R<sup>2</sup> | SE | P | R<sup>2</sup> | SE | P | R<sup>2</sup> | SE | P | R<sup>2</sup> | SE | P | R<sup>2</sup> | SE | P |
| ***g*** | 0.079 | 0.041 | 0.054 | 0.010 | 0.013 | 0.477 | 0.071 | 0.038 | 0.064 | 0.046 | 0.030 | 0.120 | 0.006<sup>#</sup> | 0.009 | 0.538 | 0.016 | 0.018 | 0.355 |
| **TBV** | 0.033 | 0.026 | 0.213 | 0.106 | 0.046 | 0.020 | 0.185 | 0.051 | **0.00025** | 0.111 | 0.045 | 0.015 | 0.098 | 0.043 | 0.023 | NA | NA | NA |
| **GMV** | 0.051 | 0.032 | 0.113 | 0.108 | 0.046 | 0.018 | 0.171 | 0.049 | **0.001** | 0.108 | 0.047 | 0.022 | 0.108 | 0.052 | 0.037 | 0.110 | 0.052 | 0.036 |
| **NAWM** | 0.014 | 0.017 | 0.403 | 0.083 | 0.041 | 0.045 | 0.170 | 0.052 | **0.001** | 0.096 | 0.042 | 0.023 | 0.094 | 0.046 | 0.041 | 0.063 | 0.034 | 0.067 |
| **WMHV** | 0.002 | 0.006 | 0.807 | 0.040 | 0.032 | 0.212 | 0.002<sup>*</sup> | 0.006 | 0.747 | 0.002 | 0.006 | 0.732 | 0.004 | 0.008 | 0.591 | 0.003 | 0.007 | 0.707 |
| ***g*FA** | 0.005 | 0.008 | 0.540 | 0.005 | 0.007 | 0.496 | 0.005 | 0.009 | 0.563 | 0.016 | 0.016 | 0.344 | 0.004 | 0.009 | 0.667 | 0.004 | 0.008 | 0.583 |
| ***g*MD** | 0.012 | 0.015 | 0.425 | 0.018 | 0.017 | 0.290 | 0.004 | 0.007 | 0.550 | 0.008 | 0.011 | 0.444 | 0.003 | 0.006 | 0.634 | 0.005 | 0.007 | 0.526 |

UPLC-MS-POS lipids together predicted 19% (SE 5%, p=0.00025) of the variance in total brain volume and 17% (SE 5%, p=0.001) of the variance in both grey matter and normal appearing white matter volumes (Table 2). Nineteen of the 193 UPLC-MS-POS lipids occurred in more than 75% of iterations for at least two of the significant models (Supplementary Table 5). These included lipids from the following lipid subclasses: sphingomyelins, ceramides, fatty acylcarnitines, monoacylglycerols, diacylglycerophosphocholines and monoacylglycerophosphocholines. Sphingomyelin, SM(d18:2/14:0), occurred in 100% of iterations of all three statistically significant models.

No set of metabolites predicted cognitive ability, WMH volume, or either measure of white matter connectivity.

Using rank-based inverse normal transformed metabolites in the models, indicated that significant predictions were not being driven by outlying metabolite measurements, as results remained essentially the same (Supplementary Table 6). Correlations between total brain, grey matter and normal appearing white matter volumes, and intracranial volume ranged from 0.8-0.9 (Supplementary Table 7). Correcting total brain, grey matter and normal appearing white matter volumes for intracranial volume greatly attenuated the results (Supplementary Table 6), indicating that the associations were mainly with absolute brain volumes and not age-related brain volume atrophy.

## Discussion

This study tested a variety of metabolite arrays for their ability to predict important cognitive and brain structure phenotypes in later life. UPLC-MS array analysis followed by penalized regression models (LASSO) identified sets of lipids that made statistically significant and substantial predictions of total brain, grey matter and normal appearing white matter volumes in the LBC1936 sample at the approximate age of 73 years.

The best performing single lipid, SM(d18:2/14:0), occurred in 100% of iterations of all three significant models, indicating a robust association which stood out beyond other lipids. SM(d18:2/14:0) is a sphingomyelin previously shown, in combination with three other compounds, to discriminate between stable and unstable angina ^55^ and to be associated with obesity ^56^. To our knowledge, it has not previously been associated with brain volume or other brain imaging related phenotypes. A second sphingomyelin, SM(d18:0/16:0), occurred in >85% of iterations of at least two of the significant models. The ceramide Cer(d18:1/16:0), which occurred in 99 or 100% of iterations of all three significant models, was previously associated with major adverse cardiovascular events ^57^. A second ceramide, Cer(d19:1/24:0), occurred in >79% of iterations of all three significant models.

Sphingolipids, including sphingomyelin and ceramide, are enriched in the central nervous system and sphingomyelin is a component of myelin sheaths. Levels of sphingomyelin and ceramide increase with age ^58–60^ and both levels and ratios of these sphingolipids have previously been associated with neurodegenerative disease ^61–63^, cognitive function, cognitive decline, and changes in white matter microstructure ^64^. Sphingomyelin degradation leads to increased ceramide ^65^. Ceramide regulates cell growth, differentiation, and apoptotic signaling in cells types, including neurons ^66^. Sphingolipids may, therefore, contribute to the prediction of brain tissue volumes in the LBC1936 via these mechanisms.

A fatty acylcarnitine [CAR(26:0)] occurred in 99 or 100% of iterations of all three significant models. Fatty acylcarnitines are important for energy production and levels are often altered in fatty acid oxidation disorders ^67^. Five other fatty acylcarnitines occurred in >75% of iterations of at least two of the significant models. Other lipid classes occurring in >75% of iterations of at least two of the significant models included, monoacylglycerols, diacylglycerophosphocholines and monoacylglycerophosphocholines, indicating that a number of lipid subclasses may be important in predicting brain volume in later-life. Although sets of lipids predict later-life brain tissue volumes in the LBC1936, this study provides no evidence that they can be used to predict later-life general cognitive ability, WMH volume or white matter connectivity. However, as we refine the lipid sets, prediction of these may be possible in the future. Future analyses could also investigate specific cognitive domains, rather than general cognitive ability, as these may be more sensitive to the effects of lipids.

Sets of lipoproteins, as measured using NMR, did not predict general cognitive ability or brain structure in this study. It may be that they are not as predictive as the lipids themselves, or that they are not as sensitive to detection by the methods used in this study.

Limitations of this study included the fact that the metabolites were measured in blood plasma rather than cerebrospinal fluid. However, plasma is a much less invasive material to collect and therefore biomarkers that can be measured in blood are particularly useful. The implications of our choice of LASSO models over other forms of penalized regression should also be borne in mind. The LASSO, compared to the related Elastic Net regression, tends to provide a sparser—and thus more readily-interpretable—set of selected features. However, given the very large number of predictors included in some of our models, the LASSO was more computationally feasible, since it does not require calculation of the additional hyperparameters required for an Elastic Net analysis. Other strengths of this study include the fact that the metabolites, cognitive ability, and structural brain MRI variables were measured at about the same time in ∼600 individuals with a narrow age range and from an ancestrally homogeneous population. Our penalized regression analysis allowed us to run prediction models that were robust to overfitting (since we split our overall sample into training and test subsets, and used cross-validation in the training subset), and multicollinearity. The form of modelling we employed to deal with the cognitive variables, latent variable structural equation modelling, reduced the impact of measurement error specific to any individual task or domain of cognitive ability, and allowed us to examine more directly each participant’s cognitive ability at a highly general level.

In conclusion, sets of lipids, particularly sphingolipids measured using UPLC-MS, predicted up to 19% of the variance in brain tissue volumes in an older relatively healthy Scottish cohort. These results suggest that future studies should concentrate on identifying specific lipids as potential cognitive and brain-structural biomarkers in older individuals.

## Supporting information

Supplementary information is available here

## Acknowledgments

We thank the LBC1936 participants and team members who contributed to these studies. Phenotype collection was supported by Age UK (The Disconnected Mind project). MRI brain imaging was supported by Medical Research Council (MRC) grants G0701120, G1001245, MR/M013111/1 and MR/R024065/1. Metabolomics were supported by the Dementia Platform UK MRC grant MR/L023784/2. This work was supported by the MRC and National Institute for Health Research [grant number MC_PC_12025] and infrastructure support was provided by the National Institute for Health Research (NIHR) Imperial Biomedical Research Centre (BRC). The work was undertaken by The University of Edinburgh Centre for Cognitive Ageing and Cognitive Epidemiology, part of the cross council Lifelong Health and Wellbeing Initiative; funding from the Biotechnology and Biological Sciences Research Council (BBSRC) and MRC is gratefully acknowledged (MR/K026992/1).We gratefully acknowledge the contribution of co-author Professor John M. Starr, who died prior to the publication of this manuscript.

## Conflict of Interest

The authors have nothing to disclose.

## Supplementary Information

Supplementary information is available here.

## Notes

### Competing Interest Statement

The authors have declared no competing interest.

